# zAMP and zAMPExplorer: Reproducible Scalable Amplicon-based Metagenomics Analysis and Visualization

**DOI:** 10.1101/2025.03.09.633768

**Authors:** Valentin Scherz, Sedreh Nassirnia, Farid Chaabane, Violeta Castelo-Szekely, Gilbert Greub, Trestan Pillonel, Claire Bertelli

## Abstract

**Summary:** To enable flexible, scalable, and reproducible microbiota profiling, we have developed zAMP, an open-source bioinformatics pipeline for the analysis of amplicon sequence data, such as 16S rRNA gene for bacteria and archaea or ITS for fungi. zAMP is complemented by two modules, one to process databases to optimize taxonomy assignment, and the second to benchmark primers, databases and classifier performances. Coupled with zAMPExplorer, an interactive R Shiny application that provides an intuitive interface for quality control, diversity analysis, and statistical testing, this complete toolbox addresses both research and clinical needs for microbiota profiling.

**Availability and Implementation:** Comprehensive documentation and tutorials are provided alongside the source code of [zAMP](https://github.com/metagenlab/zAMP) and [zAMPExplorer](https://github.com/metagenlab/zAMPExplorer) software to facilitate installation and use. zAMP is implemented as a Snakemake workflow, ensuring reproducibility by running within Singularity or Docker containers, and is also easily installable via Bioconda. The zAMPExplorer application, designed for visualization and statistical analysis, can be installed using either a Docker image or from R-universe.

## 1 Background

The microbiota is recognized as a dynamic and complex ecological community essential to environmental balance and ecosystem’s functionality. In human and veterinary medicine, increasing evidence supports its role in health and disease (Hou et al., 2022; Scherz et al., 2022). In addition to providing a biome-based understanding, microbiota profiling also offers novel possibilities for diagnostic and prognostic tools that could significantly influence clinical decision-making and patient care (Malla et al., 2019). Targeted metagenomics, or metabarcoding, by high-throughput sequencing of the *16S rRNA* encoding gene and the Internal Transcribed Spacer (ITS) regions are becoming routinely used for the characterization of specific bacterial, archaeal, and fungal communities (Armougom, 2009). It complements traditional methods used in clinical microbiology offering high sensitivity and a comprehensive detection of microbial species, particularly in samples harboring complex communities such as polymicrobial abscesses, outperforming culture methods (Gupta et al., 2019; Nam et al., 2023).

However, analyzing microbiomes remains challenging due to the diversity of sample types, the compositional complexity of microbial communities, and inconsistencies in analytical and statistical methodologies (Knight et al., 2018; Scherz et al., 2022; Trinh et al., 2018). These challenges complicate the identification of microbial taxa associated with health outcomes, interventions, or forensic investigations (Gouello et al., 2022; Trinh et al., 2018). Additionally, biases and errors in all steps from library preparation, amplification, sequencing, to bioinformatics analysis affect data accuracy and interpretability (McLaren et al., 2019; Nearing et al., 2021; Pfeifer, 2017). Notably, bioinformatics pipelines are particularly sensitive to changes in processing parameters, with even minor changes in clustering algorithms, classifiers, or database selection significantly influencing reproducibility and reliability (Kulkarni et al., 2018; Nearing et al., 2018; Siegwald et al., 2017; Ye et al., 2019). These issues underscore the importance of standardized, rigorous protocols and bioinformatic pipelines across research, clinical, and forensic applications (Fjukstad et al., 2019; Pollock et al., 2018; Scherz et al., 2022).

Commercial microbiome analysis platforms such as EzBiome (Yoon et al., 2017), llumina’s BaseSpace Sequence Hub (Illumina, 2024), One Codex (Minot et al., 2015), and QIAGEN Microbial Genomics Module (CLC Microbial Genomics Module 24.0, n.d.) offer comprehensive workflows with various features tailored for both clinical and research applications. Yet, they are paid services and require data submission to tertiary infrastructure for cloud computing, limiting accessibility for certain users (Marizzoni et al., 2020; Mbareche et al., 2020; Prodan et al., 2020; Stajich & Lapp, 2006). Open-source projects, like QIIME2 (Bolyen et al., 2019; Caporaso et al., 2010), and MOTHUR (Schloss et al., 2009) provide tutorials and tools for microbiota profiling. However, these tools can be challenging to install and use, particularly for users without bioinformatics expertise, and they may lack reproducibility and flexibility to accommodate novel workflows or specific experimental needs (Dudley & Butte, 2009; Lutz et al., 2022). Furthermore, many pipelines do not address the limitations of 16S amplicon sequencing, such as its inability to distinguish closely related taxa due to highly conserved sequences in the targeted regions (Johnson et al., 2019).

In-house bioinformatics pipelines present an attractive alternative, offering cost-free adaptable solutions that benefit from community contributions and open-source development (Fjukstad et al., 2019; Mbareche et al., 2020; Scherz et al., 2022). Workflow engines, such as Snakemake (Mölder et al., 2021) or Nextflow (DI Tommaso et al., 2017), improves these pipeline’s usability and efficiency by automating complex analyses and parallelizing processes (Caporaso et al., 2010).

In this setting, we have developed zAMP, an open-source Snakemake-based bioinformatics pipeline designed for scalability and ease of use. zAMP simplifies microbiota profiling through a single-command installation, introduces a database pre-processing step to improve taxonomic classification accuracy, and integrates a wide range of standard amplicon analysis tools. Additionally, zAMPExplorer, an R Shiny app (Jackson et al., 2021; R Core Team, 2014), streamlines data exploration and visualization to facilitate the analysis and interpretation of microbiota profiling from fundamental research to clinical applications.

## 2 Implementation of zAMP, a reproducible, scalable, amplicon-based metagenomics pipeline

zAMP offers remarkable flexibility by allowing users to customize key aspects of the pipeline to meet diverse research and clinical needs. In fact, users can set forward and reverse primer sequences, adjust minimum read length for read quality control, set denoiser and classifier tool for ASV inference and taxonomic assignment, respectively. This adaptability allows researchers to tailor the pipeline to their specific study design and data requirements. To ensure reproducibility, zAMP integrates a set of state-of-the art tools deployed via Singularity (Kurtzer et al., 2017) or Conda. Pipeline version, tool command line and logs are saved to a directory to ensure traceability. Additionally, classifier files obtained from training on a specific database are assigned a unique hash to ensure consistency and reproducibility.

zAMP accepts as input local reads in fastq format or SRA accessions to download reads via the SRA Toolkit (Leinonen et al., 2011). Next, reads containing forward or reverse primers are extracted with Cutadapt, trimmed and quality filtered with DADA2 (Callahan et al., 2016) filterAndTrim function. By default, reads passing quality control are denoised and chimeras are removed to output Amplicon Sequence Variants (ASVs). zAMP also offers an alternative denoising method with VSEARCH, which clusters sequences at 97% identity into Operational Taxonomic Units (OTUs). While the OTU approach reduces the interpretation of sequencing errors as biological variants, it cannot resolve fine-scale variations which could be clinically relevant to differentiate commensal or pathogenic strains (Callahan et al., 2016). To assign sequences a taxonomy, zAMP uses the naïve Bayesian classifier from RDP (Wang et al., 2007, Wang,Q. and Cole,J.R. 2024) by default. Alternatively, users can include classifications from DADA2 (Callahan et al., 2016), QIIME 2 (Dubois et al., 2022) and Decipher (Murali et al., 2018) in the output, to compare classifiers performance. Finally, zAMP offers the possibility to rarefy samples at a user-defined threshold, to mitigate biases resulting from differences in sampling depth (Sanders, 1968). Outputs are standardized (e.g. tab-delimited ASV/OTU tables, and BIOM files) and consolidated into a single phyloseq object (McMurdie & Holmes, 2013), facilitating data management and analysis in R.

Quality control is pivotal to ensure detection of wet-lab or sequencing failure as well as unsuitable parameters that could lead to spurious results (Scherz et al., 2022). zAMP incorporates FastQC (Andrews & others, 2019) to assess the quality of individual sample sequences and MultiQC (Ewels et al., 2016) to provide an aggregated quality report in each sequencing run. The pipeline also provides tables, bar plots and Krona plots to evaluate reads counts passing the different processing steps and explore sample composition. Rarefaction curves are provided to assess sequencing depth adequacy. With its streamlined, one-command execution, zAMP simplifies microbiome data analysis.

**Figure 1.**
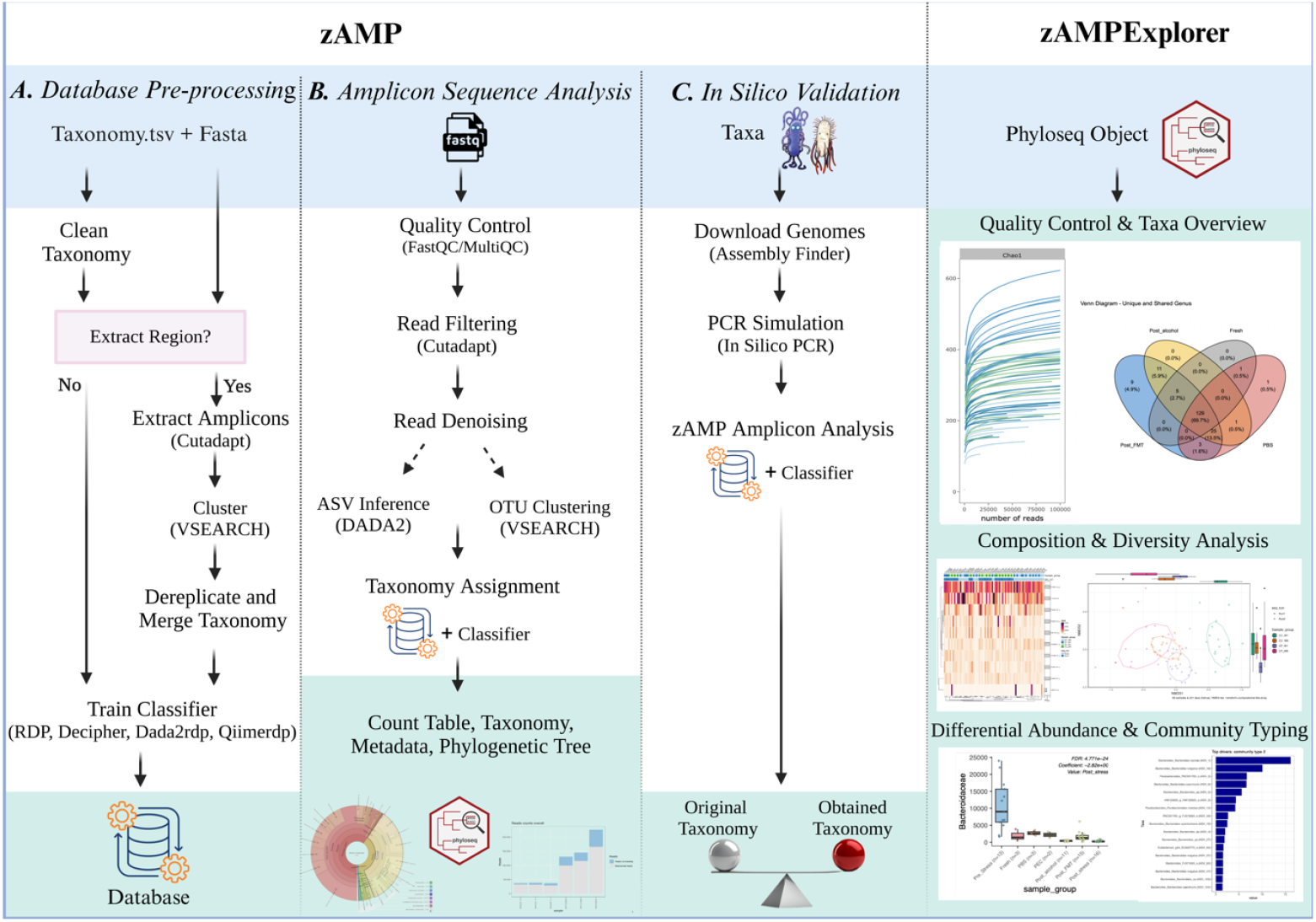
zAMP and zAMPExplorer modules. zAMP includes three modules: (A) To pre-process the reference database, and merge taxonomic ranks that cannot be distinguished based on the target region. This module has been tested with EzBioCloud, SILVA, Greengenes2, UNITE and Eukaryome databases. (B) To analyze amplicon sequences and assign taxonomic ranks. (C) To assess the classification of known genomes to evaluate the tool accuracy for specific microorganisms, given the target genomic region. zAMPExplorer, an R Shiny app, allows to visualize a phyloseq object and perform quality control and compositional analyses. The blue section represents the inputs, the green section represents the outputs, and the white section outlines the workflow and processing steps.

### 3 Secondary modules: database processing and *in silico* benchmarking

Popular 16S ribosomal RNA databases like Greengenes2 (McDonald et al., 2024) or SILVA (Quast et al., 2013) use different taxonomic nomenclatures and contain inconsistently annotated taxa (Molano et al., 2024), such as same species appearing in different genera (“convergent evolution”), duplicated names at successive ranks or empty rank names. To ensure compatibility with multiple taxonomic classifiers like RDP, zAMP searches for “convergent evolution” and raises an error when encountering such cases. Duplicated names are replaced by empty entries at their lowest rank. Empty entries are then replaced by their latest non empty name (rank propagation), in a similar approach as implemented by RESCRIPt (Robeson,M.S. et al. 2021). Currently, zAMP supports Greengenes2 (McDonald et al., 2024), SILVA (Quast et al., 2013), EzBioCloud (Yoon et al., 2017), UNITE (Abarenkov et al., 2024), and Eukaryome (Tedersoo et al., 2024) databases but it can accept any database as input provided that the fasta and taxonomy files respect the QIIME format.

As highlighted by QIIME 2’s feature-classifier plugin (Bokulich, N.A et al. 2018), tuning the parameters of taxonomic classifiers is important to improve accuracy, which includes training classifiers on the appropriate region amplified by primers. For this, zAMP allows users to process databases to extract primer amplified regions from full length sequences with Cutadapt (Martin, 2011). Extracted regions are then dereplicated with VSEARCH, and identical sequences are assigned a “merged taxonomy”. For example, *Staphylococcus aureus, schwitzerii*, and *argentus* which share an identical V3-V4 sequence will get their taxonomy merged into “*Staphylococcus aureus*/*schwitzerii/argentus”*. We hypothesized that training classifiers on the appropriate amplified region would enhance classification accuracy at species level.

zAMP pipeline incorporates an “*in silico*” module allowing to evaluate i) whether the selected PCR primers can effectively amplify specific taxa of interest, ii) the number of loci targeted by the primers on each genome, and iii) the accuracy of taxonomic classification for the resulting amplicons using different classifiers and databases. This *in silico* workflow starts by downloading genomes from NCBI using assembly_finder (Chaabane et al., 2024), before simulating PCR amplicons with simulate-PCR (Gardner & Slezak, 2014) or in-silico PCR (Ozer, 2017). Each amplicon is then assigned a taxonomy using one or multiple classifiers (RDP, qiime, DADA2…) that is compared to the expected taxonomy. To evaluate our ability to detect pathogens using the V3-V4 region of the 16S rRNA gene using different classifiers and databases, we created a mock community comprised of 1196 pathogenic bacterial species identified at the Lausanne University Hospital (Supplementary Table 1) using the *in silico* module. For Greengenes and SILVA, database processing yielded lower precision scores at all ranks, but processing the EzBioCloud database showed an improvement in classification from 0.59 to 0.74 at species level (Supplementary Fig. 1). This highlights the necessity to adapt the choice of the database, classifier and primer region according to the user needs and applications.

### 4 zAMPExplorer

zAMPExplorer is an interactive Shiny application designed to streamline and enhance microbiota analysis of amplicon-based metagenomics data. Built as a companion tool for the zAMP pipeline, it accepts a phyloseq object (the main output of zAMP) and offers a comprehensive suite of features to support each step of downstream analyses. Leveraging various R packages and libraries, zAMPExplorer automates key steps in microbiome data analysis and statistical evaluation. Starting with the upload of a phyloseq object and quality control of reads and taxa, the application enables users to explore the taxonomic composition and diversity of their samples. For deeper insights into community structure, zAMPExplorer includes community typing via Dirichlet Multinomial Modeling (DMM) (Holmes et al., 2012) and ordination visualizations like RDA plots (Dixon, 2003). With its user-friendly interface and robust functionality, zAMPExplorer empowers researchers to efficiently visualize, interpret, and analyze complex microbial data without requiring extensive coding expertise. This makes zAMPExplorer an invaluable resource for both clinical and research applications in microbiome studies.

## 5 Conclusion

To facilitate and automate microbiome analysis, from data processing to visualization and statistical evaluation, we developed zAMP, an open-access pipeline tailored for amplicon sequencing data like 16S or ITS. zAMP features a unique module for adapting and pre-processing user-defined databases for taxonomy assignment, addressing the challenges of closely related taxa. Additionally, an *in silico* validation module allows users to assess the accuracy of taxonomy assignments. Complementing the pipeline, zAMPExplorer, an interactive RShiny application, provides tools for quality control, diversity analysis, and differential abundance testing through intuitive data visualization. Together, zAMP and zAMPExplorer provide a comprehensive, reproducible, and user-friendly solution for microbiome profiling, meeting the needs of both clinical and research applications.

## 6 Acknowledgements

We would like to thank all members of the Metagenlab and other users for their feedback on the tool.

