## Supplementary Fig. 1 for "zAMP and zAMPExplorer: Reproducible Scalable Amplicon-based Metagenomics Analysis and Visualization"

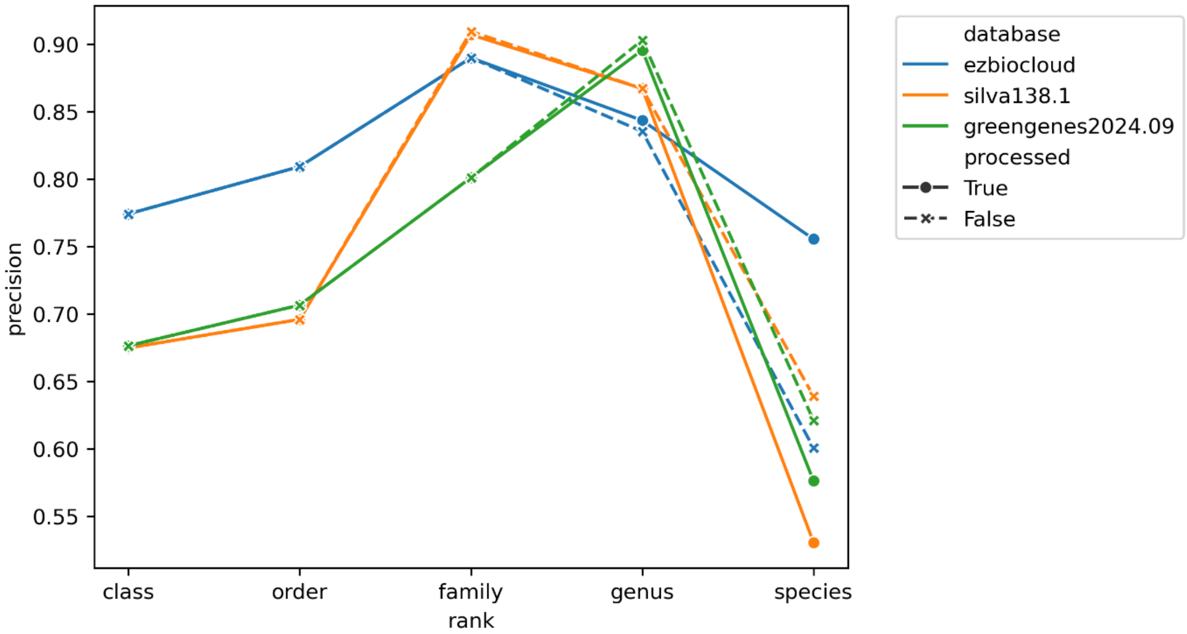


Supplementary Figure 1. Precision of sequence classification at different taxonomic levels. Three different databases EzBioCloud 2018.05, SILVA 138.1 and GreenGenes 2024.09. Of note, the publicy available version of EzBioCloud from May 2018 has been used since further releases are under licensing.

Supplementary Table 1. List of pathogens used for the benchmarking. The list consists of bacterial species identified in the diagnostic setting at Lausanne University Hospital and indicates the genome accession used for the benchmarking of the different databases using the *in silico* module.
